# Multimodal analysis identifies pericyte-centered signaling programs altered by sex and brain region in Alzheimer’s Disease

**DOI:** 10.1101/2025.08.06.668915

**Authors:** Kareem El-Ghazawi, Yonathan T. Aberra, Yashasvisai Veeramasu, Wisam A. Fares, Brant E. Isakson, Harald Sontheimer, Ukpong B. Eyo, Shayn M. Peirce

## Abstract

Pericytes are critical components of the neurovascular unit (NVU), regulating endothelial cell (EC) stability, blood-brain barrier (BBB) integrity, and neuroimmune signaling. However, their role in Alzheimer’s Disease (AD), particularly in the context of sex differences and brain region specificity, remains poorly defined. Here, we use single-nucleus RNA sequencing (snRNA-seq) to characterize transcriptional and intercellular signaling changes in pericytes across the middle temporal gyrus (MTG) and dorsolateral prefrontal cortex (DLPFC) of the same AD and non-AD donors, stratified by sex. Using LIANA and Tensor-cell2cell, we identify latent communication programs altered in female AD donors, including a pericyte-EC signaling pattern that activates TGFβ via extracellular matrix ligands and is upregulated in the MTG but not the DLPFC. A second communication pattern, downregulated in female AD donors, reveals impaired estrogen pathway signaling through ligand-receptor interactions between pericytes and astrocytes. Supporting this, we observe reduced expression of pericyte-derived neuroligins and increased pericyte-astrocyte separation in a spatial transcriptomic subset. Additionally, we identify a microglia-to-pericyte signaling program conserved across brain regions, enriched for inflammatory pathways including hypoxia and p53, and elevated in both male and female AD donors with regional specificity. This result contrasts with the more sex-and region-specific pericyte signaling programs and suggests parallel mechanisms of NVU disruption between brain regions in AD. Our findings reveal brain region-specific and sex-specific pericyte signaling changes in AD and implicate vascular-, immune-, and synapse-associated pathways in NVU dysfunction. Altogether, the data suggest pericyte-driven communication as a mechanistic contributor to female-biased vulnerability in AD and support the need for sex-aware and region-specific approaches in neurodegeneration research.

## Introduction

Alzheimer’s Disease (AD) is a progressive neurodegenerative disorder characterized by cognitive decline and widespread brain pathology, including amyloid plaques, tau tangles, neuroinflammation, and cerebrovascular dysfunction.^1^ Mounting evidence suggests that vascular pathology is not simply a consequence of AD but may actively contribute to disease onset and progression.^2,3^ In particular, the neurovascular unit (NVU)—composed of endothelial cells (ECs), pericytes, astrocytes, and microglia—plays a crucial role in maintaining blood-brain barrier (BBB) integrity, regulating cerebral blood flow, and coordinating neuroimmune interactions.^4^ Dysregulation of NVU signaling has been implicated in early AD pathology,^5^ yet the mechanisms by which specific NVU cell types contribute to these changes remain incompletely understood.

Pericytes, which enwrap capillaries and mediate communication with ECs and glia, are emerging as critical regulators of cerebrovascular health. Experimental studies have shown that pericyte dysfunction can lead to BBB breakdown,^6,7^ impaired perfusion,^8^ and altered inflammatory responses—all features observed in AD brains.^9^ However, much of our understanding of pericyte involvement in AD is derived from animal models or bulk tissue analyses, which obscure cell-type-specific and regionally localized interactions. Moreover, the extent to which pericyte signaling pathways differ between sexes remains underexplored, despite the well-established epidemiological observation that women are at greater risk for developing AD and may exhibit distinct pathological features.^10^

To address these gaps, we leveraged the Seattle AD Brain Consortium single-nucleus RNA-sequencing (snRNA-seq) data^11,12^ from the middle temporal gyrus (MTG) and dorsolateral prefrontal cortex (DLPFC) of the same postmortem human donors to construct a comprehensive, sex-stratified atlas of NVU transcriptional and intercellular signaling profiles in AD. To infer communication networks, we applied Ligand-Receptor Inference Analysis (LIANA),^13,14^ a consensus framework that integrates multiple published ligand-receptor resources and scoring methods to robustly infer cell-cell interactions based on transcriptomic data. This allowed us to compute communication scores between all NVU cell types using curated receptor-ligand pairs across a unified interface.

To uncover coordinated patterns of signaling across donors, we further applied Tensor-cell2cell,^15–17^ a dimensionality reduction framework that constructs a four-dimensional communication tensor— capturing variation across donors, ligand-receptor pairs, and sender-receiver cell-type pairs—and performs tensor decomposition to identify latent communication factors. Each factor represents a distinct pattern of cell-cell signaling activity that may underlie context-specific biological processes, such as disease state, sex, or brain region. We annotated these factors using gene set enrichment analysis (GSEA) and pathway-level footprint enrichment analysis (FEA) to gain mechanistic insight into the signaling pathways likely driving each communication program.

To evaluate the specific role of pericytes in these altered communication landscapes, we implemented a full NVU tensor decomposition in addition to a pairwise pericyte-centric tensor decomposition strategy, running LIANA + Tensor-Cell2cell separately for pericyte-astrocyte, pericyte-endothelial, and pericyte-microglial signaling axes. We then compared communication programs across brain regions and disease contexts. Additionally, we used spatial transcriptomics in a subset of donors to validate a key ligand-receptor pair that was present in both the snRNA-seq and MERFISH datasets, and its proximity relationship in situ.

Our findings reveal brain region-and sex-specific disruptions in pericyte signaling networks in AD, highlighting altered TGF-β and estrogen pathway activity as potential contributors to neurovascular dysfunction and sex-differential vulnerability. This study provides new insight into pericyte-driven communication changes in AD and underscores the importance of incorporating sex and brain region as biological variables in the study of neurodegeneration.

## Results

### A snRNA-seq dataset of the neurovascular unit from the MTG and DLPFC of the same patients

We utilized a previously published snRNA-seq provided by the Seattle Alzheimer’s Disease Brain Cell Atlas,^12^ which contains data from MTG and DLPFC of the same 84 donors (Fig. 1A). After filtering and creating subsets that only include astrocytes, pericytes, microglia, and ECs, we retained 57 donors (described in Methods) that included: females with AD (FAD), males with AD (MAD), females with no AD (FNA), and males with no AD (MNA). Group representation across sex and diagnosis was balanced (Fig. 1B–D). Cell types were confirmed by their enrichment for canonical markers (Supp Fig. 1). The total number of cells and genes between the two brain regions was also found to be similar between the two brain regions (Fig. 1C-D). We proceeded with differential gene expression and LIANA with Tensor-Cell2Cell analyses^16^ to investigate region-and sex-specific signaling alterations in the NVU. This was followed up with a pericyte-centric pairwise analysis for each of the other cell types (Fig. 1E).

**Figure 1.**
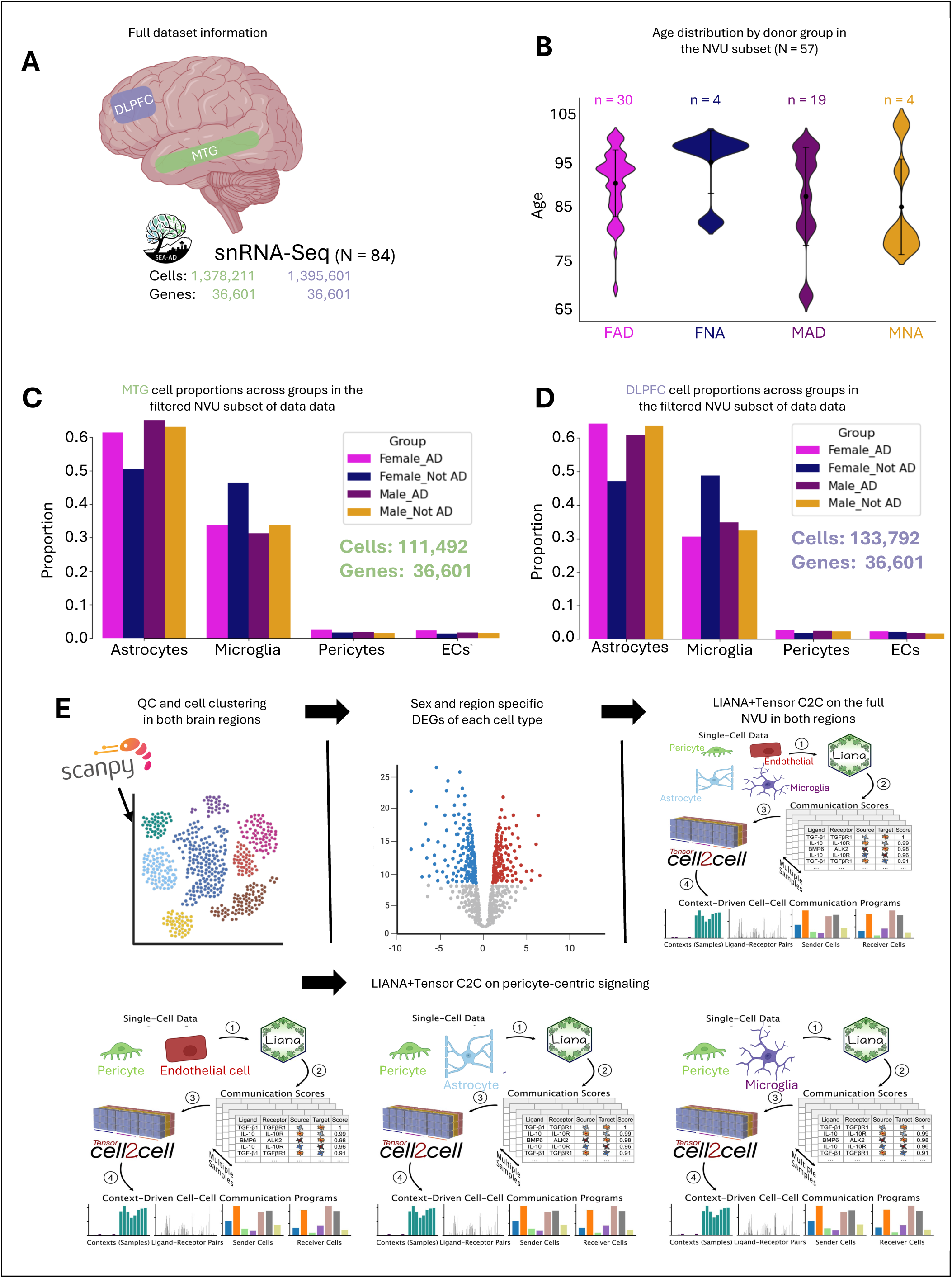
Dataset and demographic information. A) Cell and gene numbers from the full SEA-AD snRNA-Seq dataset from the MTG (green) and DLPFC (purple). B) Distribution of donor ages in each group of the NVU cell subset. C) Proportions of each NVU cell type in the MTG for each group. D) Cell proportions in the DLPFC for each group. E) Workflow of the study’s analyses adapted from Baghdassarian, Dimitrov *et al.*16

### NVU cells show a sex specific differentially expressed gene (DEG) profile

In the MTG, we identified a strong sex-specific transcriptional response. Female AD (FAD) donors showed numerous differentially expressed genes (DEGs) in pericytes, ECs, and astrocytes compared to female not-AD (FNA) donors (Fig. 2A–D), while male AD (MAD) donors showed minimal changes. Through overrepresentation analysis, upregulated genes in pericytes, ECs, and astrocytes of FAD donors were found to be enriched in oxidative phosphorylation (Supp Fig. 2) (e.g., *MT-ATP6, MT-CO3, MT-ND3*) with pericytes also showing increased cellular stress via an upregulation in *WWOX*^18^ (Fig. 2B).

**Figure 2.**
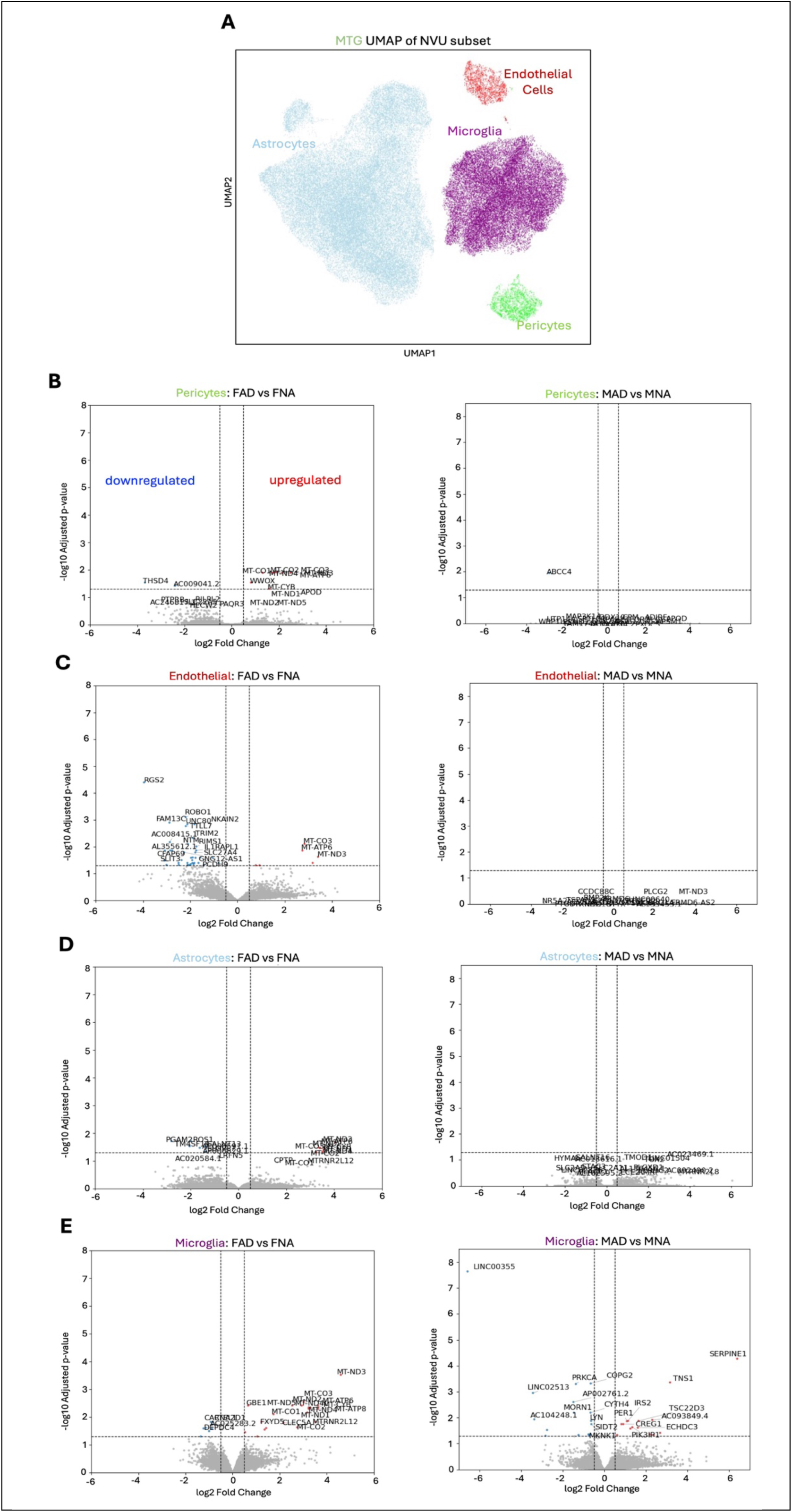
DEG analyses in the MTG. A) UMAP of all NVU cells in the MTG. B) DEGs of pericytes from FAD vs. FNA donors and MAD vs. MNA donors. C) DEGs of ECs from FAD vs. FNA donors and MAD vs. MNA donors. D) DEGs of astrocytes from FAD vs. FNA donors and MAD vs. MNA donors. E) DEGs of microglia from FAD vs. FNA donors and MAD vs. MNA donors. P-values were adjusted using the Benjamini-Hochberg method, and thresholds of adjusted-p > 0.05 and |log2FoldChange| > 0.3 were used.

ECs and astrocytes in FAD donors also showed several downregulated genes. In ECs, these downregulated genes were enriched for neurovascular/synaptic signaling (e.g. *ROBO1, NTM, SEMA5A*),^19–22^ cell adhesion (e.g. *IL1RAPL1*),^23,24^ vascular integrity (*ELN*),^25^ stress/apoptotic response (e.g. *RGS2*),^26^ cytoskeletal/structural stability (e.g. *MAP7, TRIM2*),^27,28^ and metabolism (e.g. *TAFA5, SCD5, SLC27A4*)^29–31^ (Fig. 2C). Downregulated astrocyte genes in FAD donors additionally were associated with metabolism and energy production (*PGAM2, ELOVL7*),^32,33^ cell growth and survival signaling (*ROS1*),^34^ and membrane and structural protein/cell adhesion (*TM4SF19*)^35^ (Fig. 2D). In contrast, microglia showed robust DEGs in both sexes. (Fig. 2E).

Overall, we report pericyte, EC, astrocyte, and microglial DEGs in the MTG of FAD donors but only microglial DEGs in the MTG of MAD donors. In contrast, in the DLPFC, pericytes show no DEGs in either sex, and ECs express only four DEGs across both sexes (Supp Fig. 3). These findings highlight a female-specific, MTG-localized transcriptional vulnerability in pericytes, ECs, and astrocytes in AD, suggesting that NVU dysfunction in women is associated with coordinated metabolic and structural changes across multiple vascular-associated cell types.

**Figure 3.**
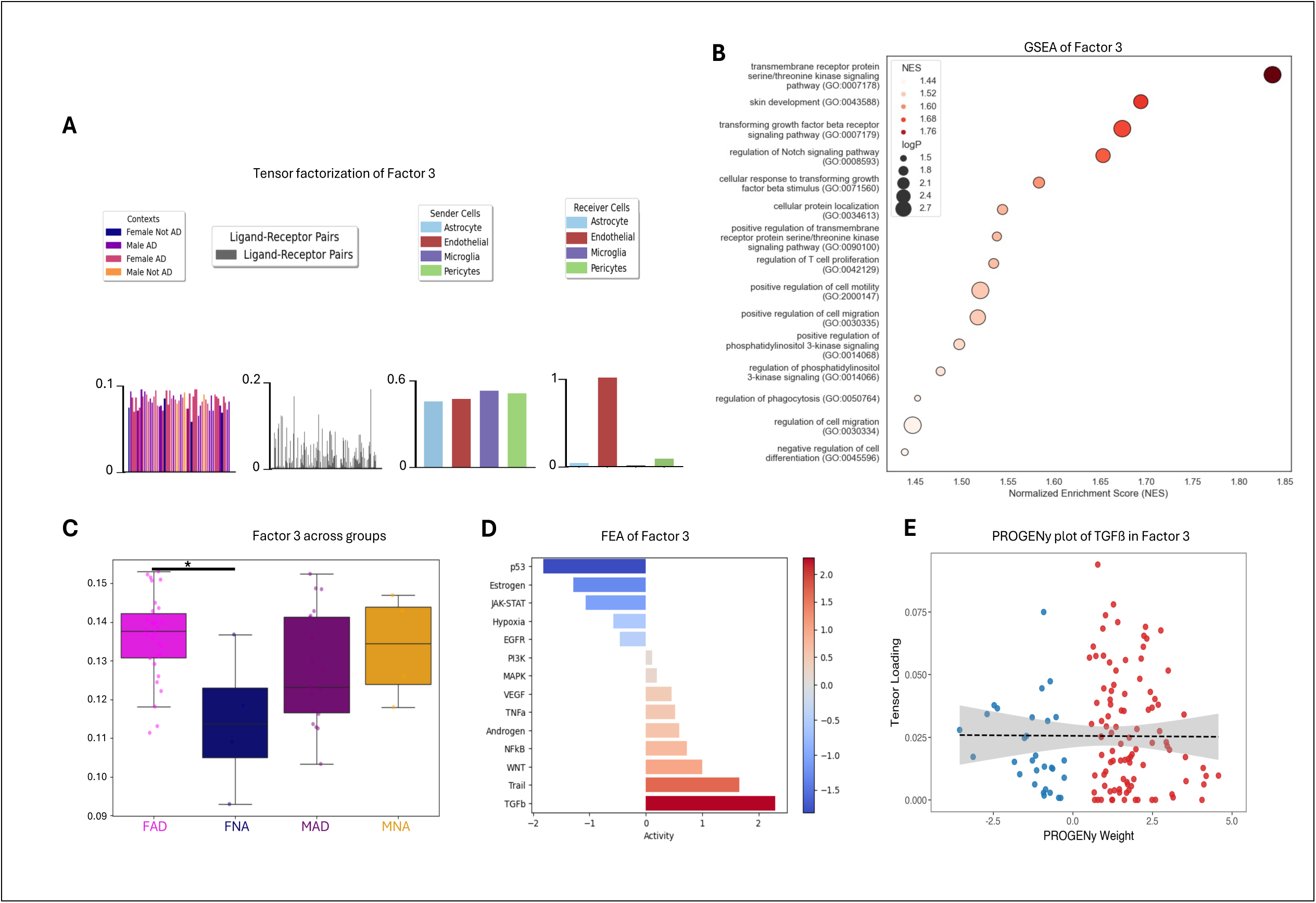
Communication factor 3 from the full NVU in the MTG across all groups. A) Tensor factorization of Factor 3. B) GSEA of Factor 3 using GO terms, NES, and log(p-value). C) Comparison of Factor 3 communication using a one-way ANOVA with Tukey’s multiple correction between all groups (*p < 0.05). D) FEA PROGENy barplot of factor 3 with active pathways in red and inactive pathways in blue. E) PROGENy plot of LR pairs contributing to TGFß activity, where each dot is a unique LR pair.

### NVU cells show a sex specific disruption of communication patterns

Tensor-cell2cell decomposition of NVU-wide ligand-receptor activity in the MTG revealed two communication programs—Factor 3 and Factor 9—that were significantly altered across sex and diagnosis groups. Factor 3 was characterized by signaling from microglia, astrocytes, and pericytes directed toward ECs (Fig. 3A). Gene set enrichment analysis (GSEA) of this factor identified strong enrichment for TGFβ signaling, cell migration, and membrane remodeling (Fig. 3B). Factor 3 activity was significantly higher in FAD donors compared to FNA donors (Fig. 3C), suggesting enhanced vascular-targeted signaling in FAD brains. To investigate pathway activity, we performed footprint enrichment analysis (FEA) using PROGENy gene sets on ligand-receptor (LR) loadings derived from Tensor-cell2cell. This analysis revealed strong TGFβ pathway activity (Fig. 3D). To pinpoint upstream ligands from sender cells that may drive this activity via EC receptors, we constructed LR-pathway gene sets based on PROGENy footprints (Fig. 3E).

This analysis highlighted several extracellular matrix (ECM)-associated ligand-receptor interactions as key contributors of TGFβ pathway activation, including *CCN2^ITGA5*, *FN1^ITGAV-ITGB1*, and *COL4A1^ITGA1-ITGB1* (Table 1).

**Table 1.**
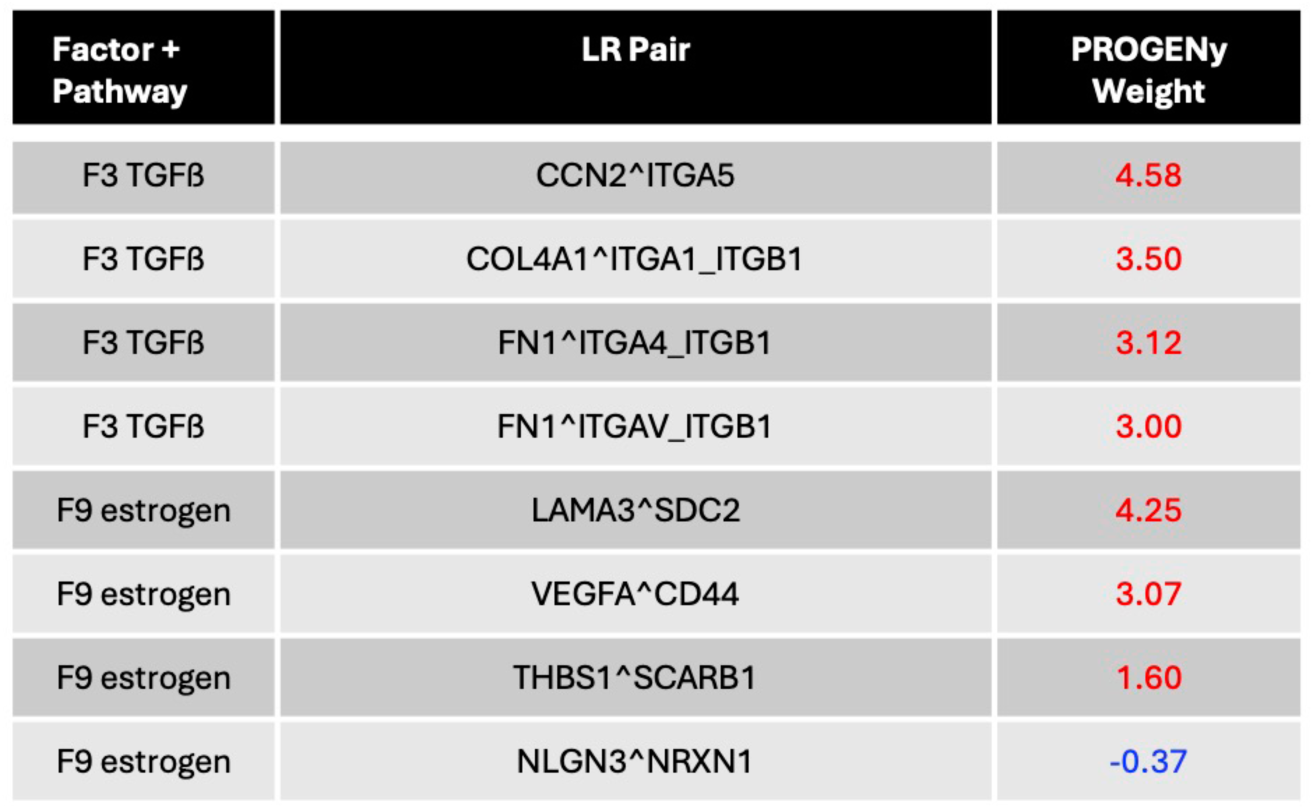
LR pairs contributing to TGFß and estrogen pathway activation in the MTG.

Factor 9 was also seen to be altered in AD. Factor 9 is involved in signaling from each of the four NVU cells to one another equally (Fig. 4A). GSEA identified this communication pattern to be involved in synapse regulation, assembly, and organization as well as cell-cell adhesion (Fig. 4B). This factor is significantly reduced in FAD donors as compared to FNA donors (Fig. 4C). FEA identified both the WNT and estrogen pathways as highly active in Factor 9 (Fig. 4D). Given the female-specific downregulation of this communication pattern, we chose to focus on the estrogen pathway due to its known roles in synaptic support, vascular integrity, and AD resilience.^36,37^ LR pathway gene set mapping revealed the estrogen signaling in this factor was shaped by both activating LR pairs involved in ECM remodeling (e.g. *LAMA3^SDC2*), angiogenesis, and immune modulation (e.g. *VEGFA^CD44 and TBS1^SCARB1*), as well as inhibitory interactions in synaptic communication (e.g *NRLGN3^NRXN1*) (Fig. 4E, Table 1). The reduction in Factor 9 in FAD donors thus reflects a loss of both estrogen-promoting and estrogen-repressing signals, suggesting a broader disruption in pericyte-associated homeostatic signaling.

**Figure 4.**
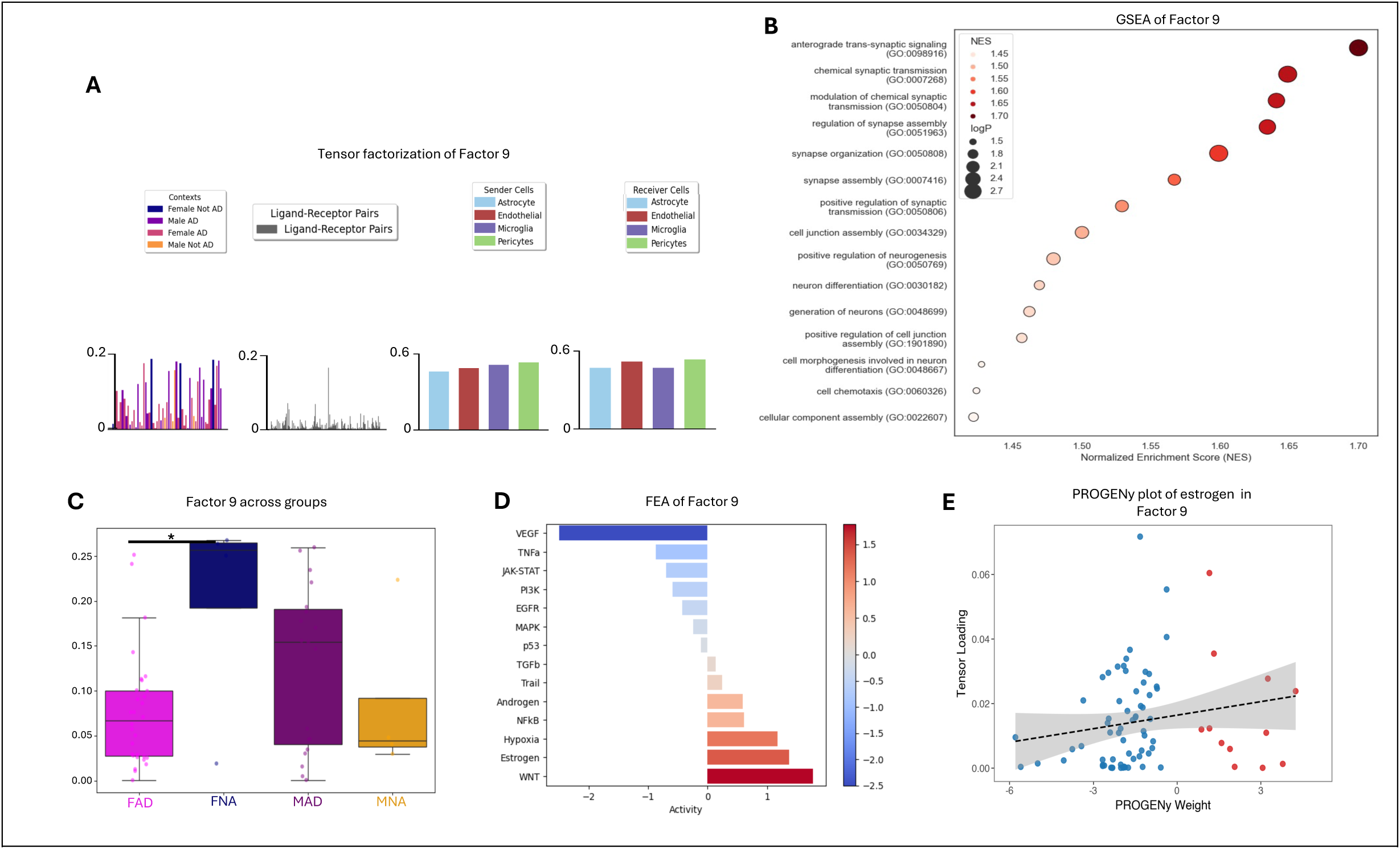
Communication factor 9 from the full NVU in the MTG across all groups A) Tensor factorization of Factor 9. B) GSEA of Factor 9 using GO terms, NES, and log(p-value). C) Comparison of Factor 9 communication using a one-way ANOVA with Tukey’s multiple correction between all groups. D) FEA PROGENy barplot of factor 9 with active pathways in red and inactive pathways in blue. E) PROGENy plot of LR pairs contributing to estrogen pathway activity, where each dot is a unique LR pair.

In the full NVU analysis of the DLPFC, no communication pattern was found to be altered between groups. More specifically, factors that were similar to Factors 3 and 9 of the MTG showed no differences between groups in the DLPFC (Supp Fig. 4). Together, these findings suggest that MTG-specific, female-enriched communication programs involving ECM-activated activated TGFβ and estrogen signaling are selectively disrupted in AD, implicating both vascular and synaptic signaling deficits in NVU dysfunction.

### Pericyte-EC signaling drives increased TGFß activity in AD relative to FNA donors

To identify how pericytes contribute to the altered communication patterns observed at the NVU level, we conducted a series of pairwise Tensor-cell2cell analyses, modeling pericyte interactions with each of the other NVU cell types in the MTG. This approach allowed us to isolate pericyte-specific signaling dynamics across cell-type pairs. Among these, the pericyte-EC interaction revealed a distinct communication program—Factor 1—that was significantly altered across disease groups. In this factor, pericytes primarily acted as signal senders to EC receivers (Fig. 5A).

**Figure 5.**
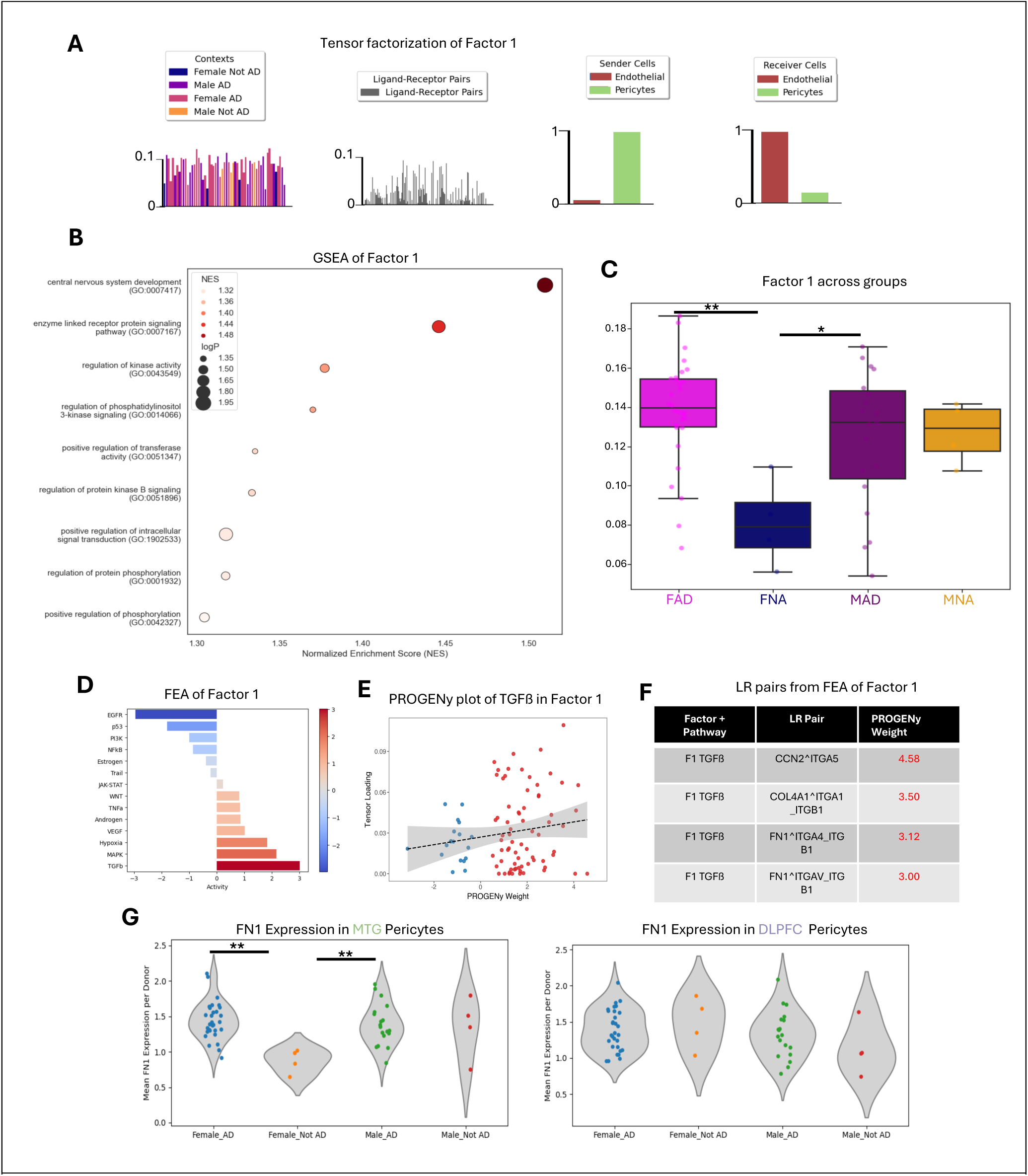
Pairwise communication analysis of pericyte-EC communication in the MTG. A) Tensor factorization of Factor 1. B) GSEA of Factor 1 using GO terms, NES, and log(p-value). C) Comparison of Factor 1 communication using a one-way ANOVA with Tukey’s multiple correction between all groups (**p <.01, *p < 0.05). D) FEA PROGENy barplot of factor 1 with active pathways in red and inactive pathways in blue. E) PROGENy plot of LR pairs contributing to TGFß activity, where each dot is a unique LR pair. F) Example LR pairs from Factor 1 with their associated weight. G) Expression of *FN1* in pericytes in each group in both MTG and DLPFC using psuedobulk, a one-way ANOVA with Tukey’s multiple correction was implemented between all groups (**p<.01).

GSEA of Factor 1 revealed enrichment for pathways associated with kinase signaling, ECM organization, and migration, mirroring the functional categories of Factor 3 from the full NVU model (Fig. 5B, see also Fig. 3B). Factor 1 activity was significantly elevated in both FAD and MAD donors compared to FNA, indicating that pericyte-driven EC signaling is upregulated in AD, with a stronger effect in females (Fig. 5C). The activated TGFß and related LR pairs also directly mirrored what we observed in Factor 3 of the full NVU analysis (Fig. 5D-F, see also Fig. 3D and Table 1). We found no such changes in the DLPFC when investigating this axis. When evaluating expression levels of the top ligands triggering TGFß activation, we found *FN1* overexpression in pericytes of both AD groups relative to FNA donors unique to the MTG (Fig. 5G). These results indicate that the increased Factor 3 activity observed at the NVU level is at least partially driven by pericyte-derived signals to ECs, highlighting a pericyte-specific contribution to vascular ECM-driven TGFβ pathway activation in the MTG that is more pronounced in FAD donors via *FN1* signaling to EC integrins.

### Pericyte-Astrocyte signaling disruption impairs estrogen activity in FAD donors

We next examined pericyte-astrocyte communication to identify additional pericyte-driven changes relevant to AD. Pairwise Tensor-cell2cell analysis between these cell types revealed a single altered communication program (Factor 9) involving bidirectional signaling between pericytes and astrocytes, as was seen in Factor 9 of the full NVU analysis (Fig. 6A). Consistent with the full NVU analysis, GSEA revealed enrichment for processes including synapse regulation, assembly, and cell-cell adhesion (Fig. 6B). Additionally, Factor 9 activity was significantly reduced in FAD donors compared to FNA donors (Fig. 6C), echoing patterns observed in the full NVU analysis (Fig. 4).

**Figure 6.**
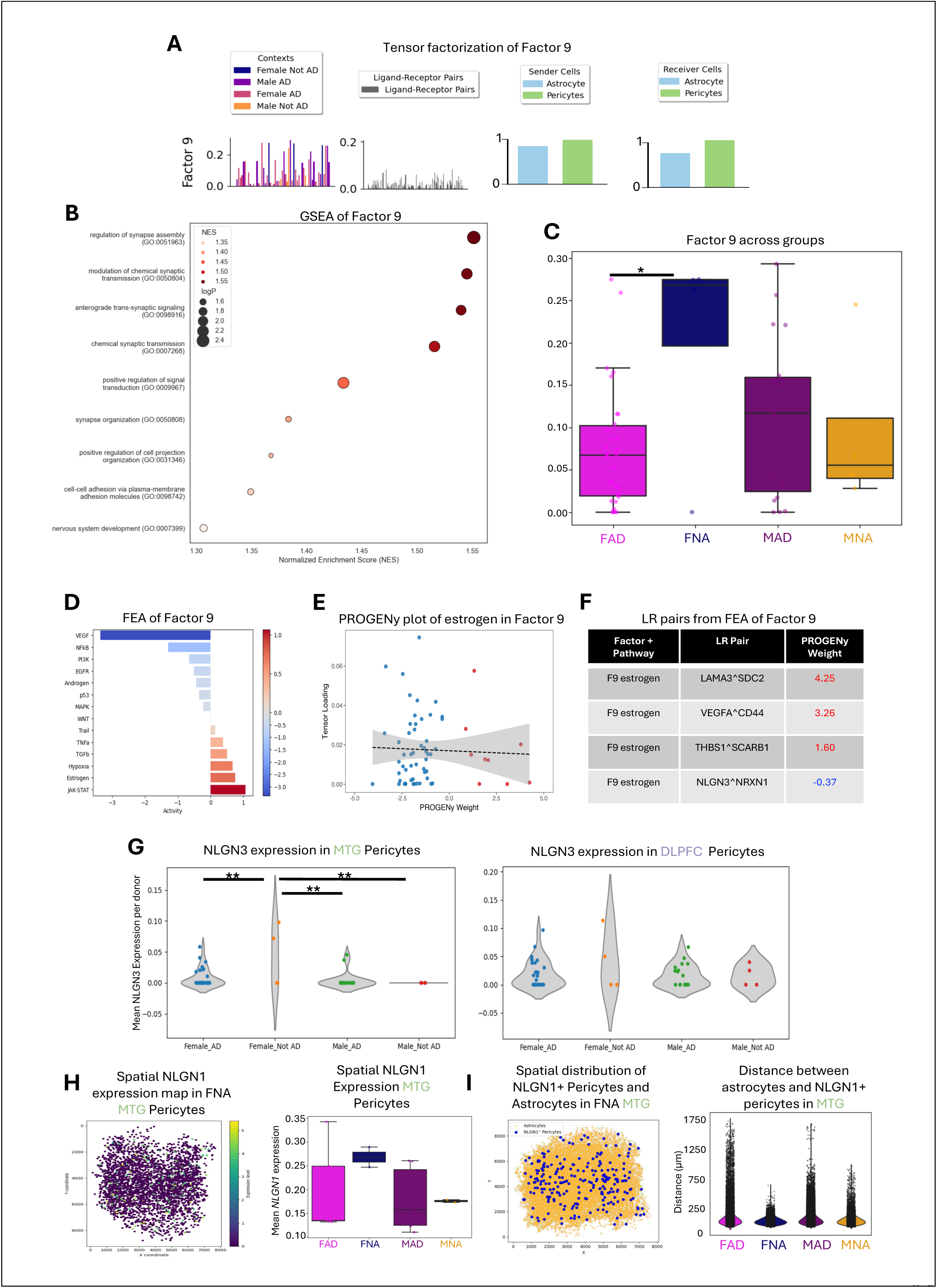
Pairwise communication analysis of pericyte-astrocyte communication in the MTG. A) Tensor factorization of Factor 9. B) GSEA of Factor 9 using GO terms, NES, and log(p-value). C) Comparison of Factor 9 communication using a one-way ANOVA with Tukey’s multiple correction between all groups (*p < 0.05). D) FEA PROGENy barplot of factor 9 with active pathways in red and inactive pathways in blue. E) PROGENy plot of LR pairs contributing to estrogen activity, where each dot is a unique LR pair. F) Example LR pairs Factor 9 with their associated weight. G) Expression of *NLGN3* in pericytes of each group in both MTG and DLPFC using psuedobulk, a one-way ANOVA with Tukey’s multiple correction was implemented between all groups (**p<.01). H) MERFISH spatial transcriptomic map of *NLGN1* expression by pericytes in the MTG of FNA donors and quantification between all groups. I) MERFISH spatial map of astrocytes and *NLGN1+* pericytes in the MTG of FNA donors and quantification between all groups.

FEA identified the estrogen signaling pathway as active in Factor 9 (Fig. 6D), with contributions from both positive regulators—such as *LAMA3^SDC2*, *VEGFA^CD44*, and *THBS1^SCARB1*—and negative regulators, including *NLGN3^NRXN1* (Fig. 6E–F). To determine whether any of these ligand-receptor interactions showed transcriptional changes consistent with the overall disruption of Factor 9 signaling, we examined the expression of all ligands and receptors contributing to estrogen pathway activity.

Among these, *NLGN3* was the only component with significant expression changes across disease groups. Notably, *NLGN3*—part of the negatively weighted *NLGN3^NRXN1* interaction—was significantly reduced in pericytes from FAD, MAD, and MNA donors relative to FNA, specifically in the MTG (Fig. 6G). While this reduction would imply decreased repression of the estrogen pathway, its presence alongside reduced overall Factor 9 activity suggests a broader collapse of pericyte-astrocyte communication balance. This pattern was not observed in the DLPFC, supporting a region-specific perturbation. Moreover, the selective expression of *NLGN3* in pericytes and *NRXN1* in astrocytes supports directional signaling from pericytes to astrocytes in this context. No changes were observed in the DLPFC.

To further explore the *NLGN3* signaling pathway, we turned to spatial transcriptomic data from a subset of the same MTG donors. Although *NLGN3* is not included in the MERFISH panel, its homolog *NLGN1* is. Like *NLGN3*, *NLGN1* is similarly reduced in all groups relative to FNA donors (Fig. 6H). Moreover, *NLGN1+* pericytes are closest in proximity to astrocytes in FNA donors as compared to other groups (Fig. 6I); confirming the reduced signaling between the cell types in those other contexts. Together, these findings support a model in which loss of pericyte-derived *NLGN3* weakens inhibitory control over estrogen signaling within Factor 9, contributing to the broader collapse of pericyte-astrocyte communication observed in the full NVU analysis. This disruption may impair estrogen-mediated synaptic maintenance mechanisms and contribute to the sex-specific vulnerability of the MTG in AD.

Given only two overlapping FNA and MNA donors had spatial transcriptomic data available, no statistical tests were performed, and these results should be interpreted with caution.

### Pericyte-Microglia communication pattern is conserved between brain regions and impaired with sex specificity

We also investigated pericyte-microglia communication to determine whether immune-related signaling contributes to disease-associated NVU changes. Pairwise Tensor-cell2cell analysis revealed a single altered communication program—Factor 2—between these cell types in the MTG, with microglia primarily acting as senders and pericytes as receivers (Fig. 7A). GSEA indicated that this factor was enriched for pathways involved in cell migration, immune activation, and inflammatory signaling (Fig. 7B). Factor 2 activity was significantly elevated in FAD donors compared to FNA (Fig. 7C), suggesting enhanced microglia-to-pericyte communication in the FAD brain. FEA identified hypoxia and p53 signaling as among the most active pathways in this factor (Fig. 7D). LR pairs activating hypoxia pathway included ECM interactions such as *ADAM17^ITGA5*, whereas p53 activation is through lipid related signaling like *APP^LRP10* and *PLTP^ABCA1* (Fig. 7E–F, Table 2).

**Figure 7.**
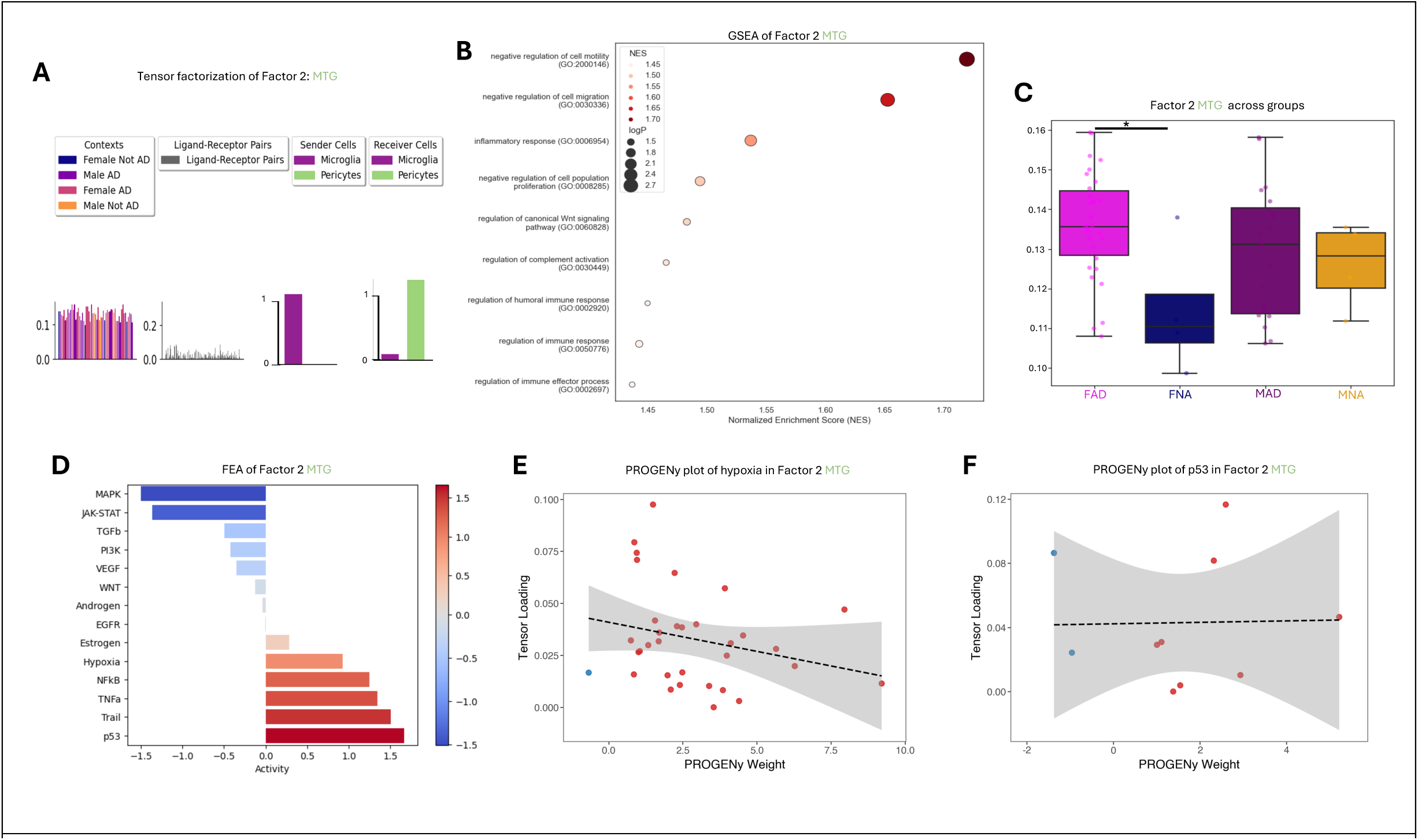
Pairwise communication analysis of pericyte-microglia communication in the MTG. A) Tensor factorization of Factor 2 of the MTG. B) GSEA of Factor 2 using GO terms, NES, and log(p-value). C) Comparison of Factor 2 communication using a one-way ANOVA with Tukey’s multiple correction between all groups (*p < 0.05). D) FEA PROGENy barplot of factor 2 with active pathways in red and inactive pathways in blue. E) PROGENy plot of LR pairs contributing to hypoxia activity, where each dot is a unique LR pair and the same for F) p53 activity. M) Example LR pairs from both factors with their associated weight.

**Table 2.**
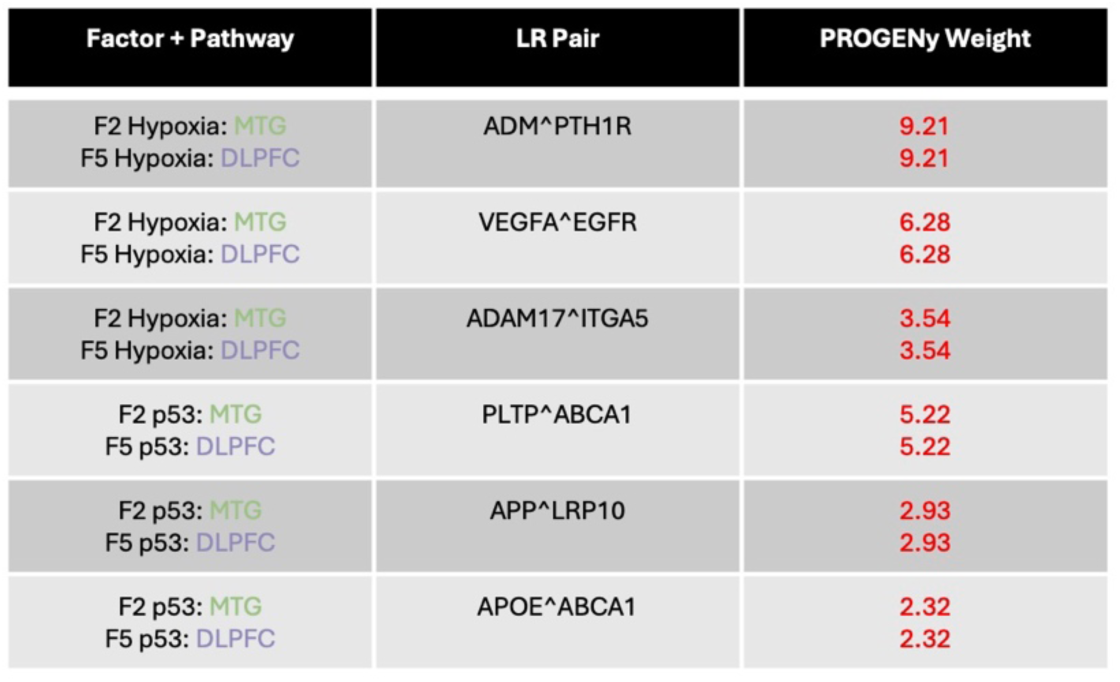
Microglia-Pericyte LR pairs activating hypoxia and p53 pathways in both MTG and DLPFC.

To evaluate whether this immune-pericyte signaling program was specific to the MTG or conserved across brain regions, we analyzed the DLPFC using the same approach. Remarkably, the same communication pattern with microglia sending to pericytes and enriched for the same inflammatory pathways—was also observed in the DLPFC (here as Factor 5) (Fig. 8A–F). However, in this region, Factor 5 was elevated in both FAD and MAD donors compared to MNA donors, indicating that the sex specificity seen in the MTG is not preserved in the DLPFC (Fig. 8C).

**Figure 8.**
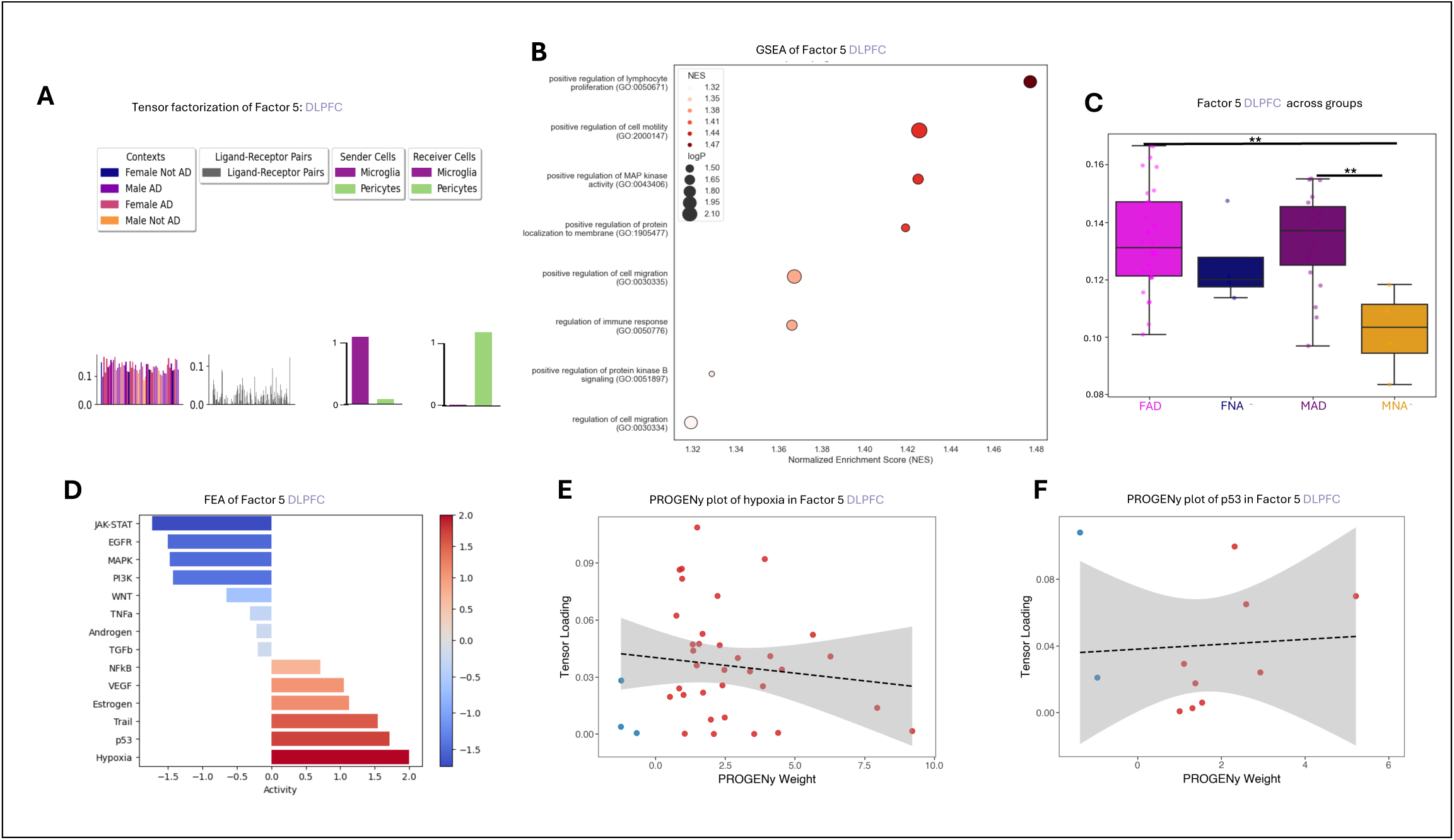
A) Tensor factorization of Factor 5 of the DLPFC. B) GSEA of Factor 5 using GO terms, NES, and log(p-value). C) Comparison of Factor 5 communication using a one-way ANOVA with Tukey’s multiple correction between all groups (**p < 0.01). D) FEA PROGENy barplot of factor 5 with active pathways in red and inactive pathways in blue. E) PROGENy plot of LR pairs contributing to Hypoxia activity, where each dot is a unique LR pair and the same for F) p53 activity.

In both regions, hypoxia and p53 signaling remained highly active, and the contributing LR interactions and weights were nearly identical between MTG and DLPFC (Fig. 8D–F, Table 2). Together, these findings suggest that microglia-driven inflammatory signaling to pericytes is a conserved feature of AD pathology across these two brain regions, but that the magnitude and sex specificity of this communication varies by brain region. These results reveal a conserved microglia-to-pericyte inflammatory signaling program—absent in the full NVU analysis but uncovered through pairwise modeling—that is enriched for hypoxia and p53 activity and may contribute to pericyte dysfunction in AD. While this communication axis is present in both MTG and DLPFC, its female specificity in the MTG and broader presence in males in the DLPFC suggest that microglia-driven pericyte signaling may contribute to sex-and region-specific patterns of neurovascular vulnerability.

## Discussion

In this study, we applied a multimodal computational framework to dissect cell-cell communication within the human NVU in AD, with a focus on sex-and region-specific differences. Using snRNA-seq data from the MTG and DLPFC of the same donors, we combined DEG analysis with ligand-receptor inference via LIANA, latent communication modeling via Tensor-cell2cell, pathway activity estimation via FEA, and spatial transcriptomics. This integrative approach enabled us to identify transcriptional and communication-level shifts that would not be evident from gene expression alone.^38^

Our DEG analysis revealed a sex-and region-specific pattern: FAD donors exhibited widespread transcriptional changes in pericytes, astrocytes, and ECs in the MTG, while MAD donors only showed changes in microglia of the MTG. In the DLPFC, changes were primarily seen in microglia with fewer in astrocytes of MAD donors. Building on these transcriptional patterns, we identified two latent intercellular communication programs of the full NVU—Factor 3 and Factor 9—that were significantly altered in the MTG of FAD donors, but not in the DLPFC.

The marked asymmetry in DEGs between female and male donors in the MTG underscores a pronounced sex-specific molecular response in AD. Although not evaluated in MTG, recent studies have also shown different transcriptional profiles between female and male AD, with female AD patients having more DEGs.^39–41^ In our findings in the MTG, pericytes, astrocytes, and ECs from FAD donors exhibited coordinated transcriptional changes, including upregulation of oxidative phosphorylation and stress-response genes and downregulation of adhesion and metabolic regulators. In contrast, these cell types in MAD donors showed minimal DEG signatures in the MTG. Previous work has similarly identified female-specific DEG profiles for EC, microglia, and astrocytes; although pericytes were not evaluated.^42^ While DEG analysis in the DLPFC revealed limited transcriptional changes overall, we did observe a set of DEGs in astrocytes from MAD donors, suggesting a cell-type and sex-specific signal in that region. This transcriptional landscape provided the foundation for the subsequent communication modeling, where altered signaling programs were found exclusively in the MTG and primarily in FAD donors.

The alignment between cell-type-specific DEG patterns and Tensor-cell2cell factor changes suggest that latent communication alterations are not artifacts of modeling but reflect underlying shifts in gene expression relevant to ligand and receptor availability. For example, the observed upregulation of ECM components such as *FN1* in FAD pericytes corresponded with the increased activity of a pericyte-EC signaling program driving TGFβ activation. Conversely, the loss of *NLGN3* expression in pericytes paralleled the reduction in an estrogen-associated pericyte-astrocyte communication pattern. These findings illustrate how transcriptional dysregulation in AD—particularly in FAD donors and in the MTG—manifests at the level of intercellular signaling and reinforces the biological relevance of stratifying analyses by both sex and brain region.

Among the altered communication programs identified in the MTG, Factor 3 was characterized by signaling from pericytes, astrocytes, and microglia to ECs and was strongly enriched for TGFβ signaling, cell migration, and ECM remodeling. These processes have been suggested as contributors to neurovascular dysfunction in AD.^43^ Importantly, this signal was selectively increased in FAD donors, suggesting that dysregulated signaling toward the vascular endothelium may be sex-specific and regionally confined. FEA further confirmed robust activation of the TGFβ pathway, pointing to ECM-integrin interactions as primary upstream drivers.

To determine the predominate cellular source of this signaling, we performed pericyte-centric Tensor-cell2cell pairwise analyses and identified a pericyte-EC communication program (Factor 1) that mirrored the functional and pathway enrichment profiles of Factor 3. This pairwise analysis confirmed that pericytes are the dominant senders, engaging EC integrins through ligands such as *FN1*, *COL4A1*, and *CCN2*. Expression analysis revealed that *FN1* was significantly upregulated in pericytes from both FAD and MAD donors, but only in the MTG, supporting a region-specific pericyte-driven mechanism of TGFβ activation. These findings position pericytes as key initiators of vascular remodeling in AD, particularly in female donors, and highlight the potential for ECM-integrin signaling axes to mediate neurovascular dysfunction via TGFß activation. ECM-integrin interactions are known to coordinate the release of TGFß from latent TGFß binding proteins in the ECM,^44^ and this ECM driven activation of TGFß has been previously described in the vascular context for Vascular Dementia,^45^ but not directly in AD or with any indication of sex and region specificity. It has also yet to be described in the context of pericyte-EC interactions in the brain.

The second major communication pattern altered in the MTG of FAD donors was Factor 9, which involved distributed signaling between all NVU cell types but was particularly enriched for pericyte-astrocyte interactions and processes related to synapse organization and cell-cell adhesion. FEA of this factor revealed activity of both the estrogen and WNT signaling pathways. Estrogen plays well-established roles in supporting synaptic plasticity, neuroprotection, and cerebrovascular integrity— functions that closely align with the processes enriched in this communication pattern.^46–49^

Interestingly, Factor 9 included both activating and inhibitory ligand-receptor pairs contributing to estrogen signaling. To prioritize the interactions most relevant to the observed reduction in Factor 9 activity in FAD donors, we examined the expression of all estrogen-associated ligands and receptors*. NLGN3^NRXN1* emerged as the only pair showing a transcriptional change consistent with the communication loss: *NLGN3* was significantly downregulated in pericytes from all disease groups compared to FNA donors, specifically in the MTG. *NLGN3* carried a negative weight in the PROGENy model—suggesting it acts as a repressor of estrogen signaling—its downregulation occurred alongside a broader reduction in Factor 9 activity and in other estrogen-associated LR pairs. This suggests a general collapse of estrogen-linked signaling, rather than a compensatory disinhibition of the pathway.

To further support this model, we leveraged spatial transcriptomic data. While *NLGN3* was not present in the MERFISH panel, we used its close homolog, *NLGN1,* as a proxy. *NLGN1* was similarly downregulated in pericytes from all disease groups relative to FNA donors, and *NLGN1+* pericytes were located closer to astrocytes in FNA donors than in AD groups. These findings provide spatial and transcriptional evidence for reduced pericyte-astrocyte neuroligin-neurexin interactions in AD. Together, these results point to a selective loss of estrogen-associated pericyte-astrocyte communication in the MTG of female AD donors, potentially contributing to sex-specific deficits in synaptic maintenance and neurovascular support. The estrogen pathway has been suggested to play a role in AD prevention,^50^ but its relation to neurovascular functions is still not well understood. Although *NLGN1* has previously been shown to be decreased in female AD models, ^51^ its specific role in pericyte-astrocyte signaling is not known, nor is the role of *NLGN3* in AD. More work is needed to understand how this signaling pathway relates to the estrogen pathway in AD.

In addition to the sex-specific communication programs identified in the MTG, our analysis uncovered a conserved inflammatory signaling axis between microglia and pericytes through Factor 2. In both the MTG and DLPFC, this factor was characterized by microglia acting as dominant senders and pericytes as receivers. GSEA and FEA linked this communication program to immune activation, cell migration, and signaling through hypoxia and p53 pathways—processes associated with vascular injury, stress response, and neuroinflammation in AD.

Although the structure of this communication pattern was conserved across regions, its magnitude and sex specificity were variable. In the MTG, Factor 2 activity was elevated specifically in FAD donors, while in the DLPFC, the same pattern (Factor 5) was elevated in both FAD and MAD donors. This variation may reflect differential microglial reactivity, regional vulnerability of the vasculature, or local environmental cues influencing immune-vascular crosstalk. Microglia have been shown to have a sex-and regional-specific transcriptional profile and injury response in the brain.^52–54^ They are also known to be in contact with pericytes,^55^ but more work is needed to understand the signaling pathways between these cells and their specificity to sex and brain region.

Both p53 and hypoxia are central microglial pathways in AD, which makes sense why they are seen to be conserved across regions.^56–60^ Conservation of these pathways across regions and contexts also reinforces the robustness of our modeling framework and highlights pericytes as key downstream targets of microglial-derived inflammatory signals, motivating avenues of future work to determine pericyte involvement with microglia in sex-and region-specific facets.

Collectively, our findings reveal that pericyte-centered communication programs in the MTG are selectively disrupted in FAD donors. These include a loss of estrogen-linked pericyte-astrocyte signaling, enhanced ECM-driven pericyte-EC TGFβ activation and increased immune cell-to-pericyte signaling from microglia. While estrogen pathway disruption may contribute to synaptic vulnerability in women, the enhanced TGFβ signaling may reflect a parallel sex-biased mechanism of vascular remodeling. Together, these communication programs may help explain the greater burden of neurovascular and cognitive decline observed in women.^61^ An additionally interesting observation was the consistently observed difference in healthy males and females. Although not statistically significant in their comparison, their trend in differing from one another opens an avenue for further investigation into sex-specific vulnerabilities to AD associated with aging.

Our results also emphasize the importance of regional stratification in neurodegenerative research. The MTG emerged as a uniquely affected region, with both transcriptional and cell-cell communication changes that were not observed in the DLPFC. While the DLPFC is commonly included in transcriptomic studies of AD,^62–65^ our data suggest that the MTG may be more sensitive to NVU dysfunction, especially in females. These findings support the need for future studies to more deeply investigate the MTG, particularly in relation to sex-specific pathways, neurovascular health, and synaptic integrity. More broadly, our work underscores the value of integrating single-cell, communication modeling, and spatial approaches to resolve subtle but disease-relevant signaling alterations that span both transcriptional and intercellular scales. This work can be expanded with the use of experimental validations and with computational tissue-level modeling, as our group has previously performed in other disease conditions,^66,67^ to further identify key contextual contributors in AD.

While our analysis provides new insights into context-specific neurovascular signaling in AD, several limitations should be acknowledged. First, our inference of cell-cell communication is based on transcriptomic data and curated ligand-receptor databases, which do not account for post-transcriptional regulation, protein abundance, or ligand activation states. Consequently, predicted interactions may not fully represent functional signaling. Second, although we applied both pairwise and full NVU models, these remain computational and do not capture spatial proximity or biophysical constraints *in vivo*.

Third, the analysis is based on post-mortem human brain tissue from the MTG and DLPFC, representing a static snapshot of signaling and limiting generalizability to other brain regions or disease stages.

Moreover, while donor-matched samples help control for inter-individual variability, sampling remains limited in size and diversity. Finally, while we observed strong convergence between pericyte-centric and broader NVU signaling programs, future studies integrating spatial transcriptomics, proteomics, and functional validation will be critical to confirm the directionality and functional consequences of these interactions.

This study integrates snRNA-seq differential expression, ligand–receptor inference, latent communication modeling, and spatial transcriptomics to uncover sex-and region-specific neurovascular signaling changes in AD. It identifies pericytes as central mediators of vascular remodeling and synaptic vulnerability in the female MTG through TGFβ activation and estrogen-linked signaling loss. These findings underscore the importance of stratifying analyses by brain region and sex to resolve context-specific mechanisms of neurovascular dysfunction in AD.

## Methods

### Donor samples and dataset description

We analyzed publicly available single-nucleus RNA-sequencing (snRNA-seq) data from the Seattle Alzheimer’s Disease Brain Cell Atlas, which includes matched samples from the middle temporal gyrus (MTG) and dorsolateral prefrontal cortex (DLPFC) of 84 postmortem human donors. We focused on a subset of 57 donors who passed quality control and for whom all relevant metadata (sex, diagnosis, brain region) were available. Donors were originally classified based on Alzheimer’s Disease Neuropathologic Change (ADNC) scores into four categories: No, Low, Intermediate, and High. We excluded donors with Low ADNC and combined Intermediate and High ADNC groups into a single AD group, while retaining the No ADNC group as Not AD. Final groups were then constructed based on sex and AD status, resulting in four groups: female AD (FAD), female not AD (FNA), male AD (MAD), and male not AD (MNA). A subset of these donors (N = 16) also had matched spatial transcriptomic data from the MTG using MERFISH, including 2 FNA, 2 MNA, 5 FAD and 7 MAD donors.

### snRNAseq preprocessing and cell type filtering

We performed preprocessing using the Scanpy Python package (v1.10.2). We retained only astrocytes, pericytes, microglia, and ECs based on annotations provided in the original dataset. Marker gene expression across cell types was used to confirm the annotations. Donors with less than four of any cell type were removed. Cells were filtered using standard quality control metrics, removing those with fewer than 200 detected genes or more than 15% mitochondrial content. Doublets were removed by removing cells with a high number of unique molecular identifiers (> 5,500). Data were normalized using total-count normalization and log1p transformation. Batch effect was accounted for by removing batches contained less than two cells. Highly variable genes were identified to guide dimensionality reduction and visualization, but all genes were retained for differential expression and communication analyses.

### Differential gene expression analysis

Pseudobulk expression profiles were generated for each donor, brain region, and cell type by summing raw gene counts across cells. Differential expression analysis was performed using PyDESeq2 (v0.4.12), comparing FAD vs. FNA and MAD vs. MNA within each brain region. Wald tests were used to assess statistical significance of model coefficients, and p-values were adjusted using the Benjamini-Hochberg method. Genes were considered differentially expressed if they met a threshold of adjusted p < 0.05 and absolute log2 fold-change > 0.3. GSEA was used to annotate differentially expressed genes using Gene Ontology (GO) Biological Process terms.

### Cell-Cell communication inference: LIANA + Tensor cell2cell

To infer cell-cell communication programs, we used log-normalized expression with LIANA (v1.4.0) with the consensus scoring method and the Omnipath ligand-receptor resource. The resulting interaction scores were formatted into a 4D tensor (samples × ligand-receptor pairs × sender cell types × receiver cell types) and decomposed using Tensor-cell2cell (v0.7.4) to extract latent communication factors as previously described.^16^ Separate tensor decompositions were performed for the MTG and DLPFC using the full NVU cell set.

### Gene set enrichment and footprint enrichment analyses

We used GSEA (via gseapy v1.0.3) to annotate communication factors with GO Biological Process terms. To identify signaling pathway activity, we performed FEA using PROGENy gene sets. Ligand-receptor pairs were weighted by their loadings in each factor, and enrichment was assessed using a weighted Kolmogorov-Smirnov test with 1000 permutations. Pathways with adjusted p < 0.05 were considered significantly enriched.

### Pairwise pericyte-centric Tensor cell2cell analysis

To evaluate pericyte-specific signaling axes, we created three pairwise subsets of the SEA-AD dataset, each containing pericytes and one of the other NVU cell types (astrocytes, ECs, or microglia). For each pairwise subset, we ran the same LIANA + Tensor-cell2cell pipeline as was used in the full NVU analysis. Log-normalized expression was used as input for LIANA, followed by formatting of interaction scores into a 4D tensor (samples × ligand-receptor pairs × sender × receiver). Tensor decomposition was then applied to extract latent communication patterns. Factor annotations were made using GSEA and FEA and compared with NVU-level factors by correlating LR loading vectors and overlapping enriched pathways.

### Spatial transcriptomic analysis

We analyzed spatial transcriptomic data from the same donors using MERFISH measurements of the MTG. Only genes included in the MERFISH panel (180 genes) were used, and *NLGN3* was not among them. Instead, *NLGN1* was used as a proxy. Gene expression was normalized to 10,000 counts per cell and log-transformed prior to group-level comparisons. To assess pericyte-astrocyte spatial relationships, we calculated the distance between each astrocyte and its nearest *NLGN1*+ pericyte using spatial transcriptomics data from the MTG. *NLGN1*+ pericytes were defined as pericytes with nonzero expression of *NLGN1*. Spatial coordinates were extracted from the MERFISH dataset, and distances were computed using Euclidean distance via a k-d tree implementation (scipy.spatial.cKDTree). For each astrocyte, the shortest distance to the nearest *NLGN1*+ pericyte was recorded. Group-wise comparisons were visualized using violin and strip plots. Statistical testing was not performed, as MNA and FNA groups contained only 2 donors each.

### Statistical analysis and visualizations

All statistical analyses were conducted using Python packages, including scanpy, scipy, and statsmodels. Visualizations were generated using matplotlib, seaborn, and plotnine. Significance was defined as p < 0.05 unless otherwise indicated. All enrichment p-values were corrected for multiple testing using the Benjamini-Hochberg method. Post hoc pairwise comparisons were performed using Dunn’s test when applicable.

## Acknowledgements

We would like to acknowledge and thank the donors who are a part of SEA-AD consortium. We thank the Allen Brain Map Community Forum for help with data access and interpretation. Funding sources included: UVA Brain Institute Pilot Transformative Neuroscience Grant (K.E, S.M.P, U.B.E), HL137112 (BEI), and HL165143 (BEI).

## Author contributions

K.E.: conceived the project, deigned the study, wrote the code, conducted analyses, interpreted analyses, created visualizations, wrote the initial draft, and reviewed and edited subsequent drafts. Y.T.A.: contributed to the initial code, contributed to study design, and revised the initial draft. Y.V. and W.A.F.: contributed to interpretation of analyses, contributed to the initial draft, and revised the initial draft. B.E.I. and H.S.: revised the initial draft. U.B.E. and S.M.P: supervision, contributed to study design, contributed to visualization design, and edited the original draft.

## References

1. Iadecola, C. & Gottesman, R. F. Cerebrovascular alterations in Alzheimer’s disease: incidental or pathogenic? Circ. Res. 123, 406–408 (2018).

2. Strickland, S. Blood will out: vascular contributions to Alzheimer’s disease. J. Clin. Invest. 128, 556–563 (2018).

3. Eisenmenger, L. B. et al. Vascular contributions to Alzheimer’s disease. Transl. Res. J. Lab. Clin. Med. 254, 41–53 (2023).

4. Schaeffer, S. & Iadecola, C. Revisiting the neurovascular unit. Nat. Neurosci. 24, 1198–1209 (2021).

5. Gerrits, E. et al. Neurovascular dysfunction in GRN-associated frontotemporal dementia identified by single-nucleus RNA sequencing of human cerebral cortex. Nat. Neurosci. 25, 1034–1048 (2022).

6. Benarroch, E. What Are the Roles of Pericytes in the Neurovascular Unit and Its Disorders? Neurology 100, 970–977 (2023).

7. Li, M. et al. Microvascular and cellular dysfunctions in Alzheimer’s disease: an integrative analysis perspective. Sci. Rep. 14, 20944 (2024).

8. Hartmann, D. A. et al. Brain capillary pericytes exert a substantial but slow influence on blood flow. Nat. Neurosci. 24, 633–645 (2021).

9. El-Ghazawi, K., Eyo, U. B. & Peirce, S. M. Brain Microvascular Pericyte Pathology Linking Alzheimer’s Disease to Diabetes. Microcirculation 31, e12877 (2024).

10. Moutinho, S. Women twice as likely to develop Alzheimer’s disease as men — but scientists do not know why. Nat. Med. 31, 704–707 (2025).

11. Seattle Alzheimer’s Disease Brain Cell Atlas (SEA-AD) - Registry of Open Data on AWS. https://registry.opendata.aws/allen-sea-ad-atlas/.

12. Gabitto, M. I., et al. Integrated multimodal cell atlas of Alzheimer’s disease. Preprint at 10.1101/2023.05.08.539485 (2024).

13. LIANA: a LIgand-receptor ANalysis frAmework. https://saezlab.github.io/liana/.

14. Dimitrov, D. et al. LIANA+ provides an all-in-one framework for cell–cell communication inference. Nat. Cell Biol. 26, 1613–1622 (2024).

15. Armingol, E. et al. Context-aware deconvolution of cell–cell communication with Tensor-cell2cell. Nat. Commun. 13, 3665 (2022).

16. Baghdassarian, H. M., Dimitrov, D., Armingol, E., Saez-Rodriguez, J. & Lewis, N. E. Combining LIANA and Tensor-cell2cell to decipher cell-cell communication across multiple samples. *Cell Rep*. Methods 4, (2024).

17. Context Factorisation with tensor-cell2cell. https://saezlab.github.io/liana/articles/liana_cc2tensor.html.

18. Liu, S.-Y., Chiang, M.-F. & Chen, Y.-J. Role of WW domain proteins WWOX in development, prognosis, and treatment response of glioma. Exp. Biol. Med. 240, 315–323 (2015).

19. Gangaraju, S. et al. Cerebral endothelial expression of Robo1 affects brain infiltration of polymorphonuclear neutrophils during mouse stroke recovery. Neurobiol. Dis. 54, 24–31 (2013).

20. Venkannagari, H. et al. Highly Conserved Molecular Features in IgLONs Contrast Their Distinct Structural and Biological Outcomes. J. Mol. Biol. 432, 5287–5303 (2020).

21. Duan, Y. et al. Semaphorin 5A inhibits synaptogenesis in early postnatal-and adult-born hippocampal dentate granule cells. eLife 3, e04390 (2014).

22. Sadanandam, A., Rosenbaugh, E. G., Singh, S., Varney, M. & Singh, R. K. Semaphorin 5A promotes angiogenesis by increasing endothelial cell proliferation, migration, and decreasing apoptosis. Microvasc. Res. 79, 1–9 (2010).

23. Il1rapl1 interleukin 1 receptor accessory protein-like 1 [Mus musculus (house mouse)] - Gene - NCBI. https://www.ncbi.nlm.nih.gov/gene/331461.

24. Valnegri, P. et al. The X-linked intellectual disability protein IL1RAPL1 regulates excitatory synapse formation by binding PTPδ and RhoGAP2. Hum. Mol. Genet. 20, 4797–4809 (2011).

25. Duca, L. et al. Matrix ageing and vascular impacts: focus on elastin fragmentation. Cardiovasc. Res. 110, 298–308 (2016).

26. Wang, C.-H. (Jenny) & Chidiac, P. RGS2 promotes the translation of stress-associated proteins ATF4 and CHOP via its eIF2B-inhibitory domain. Cell. Signal. 59, 163–170 (2019).

27. Adler, A. et al. A structural and dynamic visualization of the interaction between MAP7 and microtubules. Nat. Commun. 15, 1948 (2024).

28. Ylikallio, E. et al. Deficiency of the E3 ubiquitin ligase TRIM2 in early-onset axonal neuropathy. Hum. Mol. Genet. 22, 2975–2983 (2013).

29. Wesołek-Leszczyńska, A., Pastusiak, K., Bogdański, P. & Szulińska, M. Can Adipokine FAM19A5 Be a Biomarker of Metabolic Disorders? Diabetes Metab. Syndr. Obes. 17, 1651–1666 (2024).

30. Zhang, Q., Sun, S., Zhang, Y., Wang, X. & Li, Q. Identification of Scd5 as a functional regulator of visceral fat deposition and distribution. iScience 25, 103916 (2022).

31. Li, Z. et al. SLC27A4-mediated selective uptake of mono-unsaturated fatty acids promotes ferroptosis defense in hepatocellular carcinoma. Free Radic. Biol. Med. 201, 41–54 (2023).

32. Li, Z. et al. Targeting the Metabolic Enzyme PGAM2 Overcomes Enzalutamide Resistance in Castration-Resistant Prostate Cancer by Inhibiting BCL2 Signaling. Cancer Res. 83, 3753–3766 (2023).

33. Wang, X., Yu, H., Gao, R., Liu, M. & Xie, W. A comprehensive review of the family of very-long-chain fatty acid elongases: structure, function, and implications in physiology and pathology. Eur. J. Med. Res. 28, 532 (2023).

34. Acquaviva, J., Wong, R. & Charest, A. The multifaceted roles of the receptor tyrosine kinase ROS in development and cancer. Biochim. Biophys. Acta BBA - Rev. Cancer 1795, 37–52 (2009).

35. Ding, L. et al. TM4SF19 aggravates LPS-induced attenuation of vascular endothelial cell adherens junctions by suppressing VE-cadherin expression. Biochem. Biophys. Res. Commun. 533, 1204– 1211 (2020).

36. Zhao, L., Woody, S. K. & Chhibber, A. Estrogen receptor β in Alzheimer’s disease: from mechanisms to therapeutics. Ageing Res. Rev. 24, 178–190 (2015).

37. Jett, S. et al. Endogenous and Exogenous Estrogen Exposures: How Women’s Reproductive Health Can Drive Brain Aging and Inform Alzheimer’s Prevention. Front. Aging Neurosci. 14, (2022).

38. Su, J. et al. Cell–cell communication: new insights and clinical implications. Signal Transduct. Target. Ther. 9, 196 (2024).

39. National Academies of Sciences, E. et al. Transcriptomic Evidence for Sex Differences in Neurodevelopmental and Neurodegenerative Disorders. in Sex Differences in Brain Disorders: Emerging Transcriptomic Evidence: Proceedings of a Workshop (National Academies Press (US), 2021).

40. Soudy, M., Bars, S. L. & Glaab, E. Sex-dependent molecular landscape of Alzheimer’s disease revealed by large-scale single-cell transcriptomics. Alzheimers Dement. 21, e14476 (2025).

41. Belonwu, S. A. et al. Sex-Stratified Single-Cell RNA-Seq Analysis Identifies Sex-Specific and Cell Type-Specific Transcriptional Responses in Alzheimer’s Disease Across Two Brain Regions. Mol. Neurobiol. 59, 276–293 (2022).

42. Sanfilippo, C. et al. Middle-aged healthy women and Alzheimer’s disease patients present an overlapping of brain cell transcriptional profile. Neuroscience 406, 333–344 (2019).

43. Grammas, P. & Ovase, R. Cerebrovascular Transforming Growth Factor-β Contributes to Inflammation in the Alzheimer’s Disease Brain. Am. J. Pathol. 160, 1583–1587 (2002).

44. Saharinen, J., Hyytiäinen, M., Taipale, J. & Keski-Oja, J. Latent transforming growth factor-beta binding proteins (LTBPs)--structural extracellular matrix proteins for targeting TGF-beta action. Cytokine Growth Factor Rev. 10, 99–117 (1999).

45. Kandasamy, M. et al. TGF-β Signaling: A Therapeutic Target to Reinstate Regenerative Plasticity in Vascular Dementia? Aging Dis. 11, 828–850 (2020).

46. Russell, J. K., Jones, C. K. & Newhouse, P. A. The Role of Estrogen in Brain and Cognitive Aging. Neurotherapeutics 16, 649–665 (2019).

47. Zárate, S., Stevnsner, T. & Gredilla, R. Role of Estrogen and Other Sex Hormones in Brain Aging. Neuroprotection and DNA Repair. Front. Aging Neurosci. 9, (2017).

48. Sheppard, P. A. S., Choleris, E. & Galea, L. A. M. Structural plasticity of the hippocampus in response to estrogens in female rodents. Mol. Brain 12, 22 (2019).

49. Bustamante-Barrientos, F. A. et al. The Impact of Estrogen and Estrogen-Like Molecules in Neurogenesis and Neurodegeneration: Beneficial or Harmful? Front. Cell. Neurosci. 15, (2021).

50. Wharton, W. et al. Potential role of estrogen in the pathobiology and prevention of Alzheimer’s disease. Am. J. Transl. Res. 1, 131–147 (2009).

51. Dufort-Gervais, J. et al. Neuroligin-1 is altered in the hippocampus of Alzheimer’s disease patients and mouse models, and modulates the toxicity of amyloid-beta oligomers. Sci. Rep. 10, 6956 (2020).

52. Brandi, E. et al. Brain region-specific microglial and astrocytic activation in response to systemic lipopolysaccharides exposure. Front. Aging Neurosci. 14, (2022).

53. Grabert, K. et al. Microglial brain region-dependent diversity and selective regional sensitivities to ageing. Nat. Neurosci. 19, 504–516 (2016).

54. Barko, K. et al. Brain region-and sex-specific transcriptional profiles of microglia. Front. Psychiatry 13, 945548 (2022).

55. Morris, G. P. et al. Microglia directly associate with pericytes in the central nervous system. Glia 71, 1847–1869 (2023).

56. Davenport, C. M., Sevastou, I. G., Hooper, C. & Pocock, J. M. Inhibiting p53 pathways in microglia attenuates microglial-evoked neurotoxicity following exposure to Alzheimer peptides. J. Neurochem. 112, 552–563 (2010).

57. Aloi, M. S., Su, W. & Garden, G. A. The p53 Transcriptional Network Influences Microglia Behavior and Neuroinflammation. Crit. Rev. Immunol. 35, 401–415 (2015).

58. Jebelli, J., Hooper, C. & Pocock, J. M. Microglial p53 activation is detrimental to neuronal synapses during activation-induced inflammation: Implications for neurodegeneration. Neurosci. Lett. 583, 92–97 (2014).

59. Butturini, E., Boriero, D., Carcereri de Prati, A. & Mariotto, S. STAT1 drives M1 microglia activation and neuroinflammation under hypoxia. Arch. Biochem. Biophys. 669, 22–30 (2019).

60. March-Diaz, R. et al. Hypoxia compromises the mitochondrial metabolism of Alzheimer’s disease microglia via HIF1. *Nat*. Aging 1, 385–399 (2021).

61. Levine, D. A. et al. Sex Differences in Cognitive Decline Among US Adults. *JAMA Netw*. Open 4, e210169 (2021).

62. Mathys, H. et al. Single-cell multiregion dissection of Alzheimer’s disease. Nature 632, 858–868 (2024).

63. Mathys, H. et al. Single-cell atlas reveals correlates of high cognitive function, dementia, and resilience to Alzheimer’s disease pathology. Cell 186, 4365–4385.e27 (2023).

64. Olayinka, O. A. et al. Single Nucleus RNA Sequencing Reveals Interaction Between Microglia and Endothelial Cells in Mixed Alzheimer’s Disease and Vascular Pathology. Alzheimers Dement. 20, e089628 (2025).

65. Wang, X. et al. Deciphering cellular transcriptional alterations in Alzheimer’s disease brains. Mol. Neurodegener. 15, 38 (2020).

66. Leonard-Duke, J. et al. Multiscale computational model predicts how environmental changes and treatments affect microvascular remodeling in fibrotic disease. PNAS Nexus 4, pgae551 (2025).

67. Rikard, S. M. et al. Mathematical Model Predicts that Acceleration of Diabetic Wound Healing is Dependent on Spatial Distribution of VEGF-A mRNA (AZD8601). Cell. Mol. Bioeng. 14, 321–338 (2021).

