## Supplementary figures and images for "Multimodal analysis identifies pericyte-centered signaling programs altered by sex and brain region in Alzheimer’s Disease"

### Supplemental Figure 1

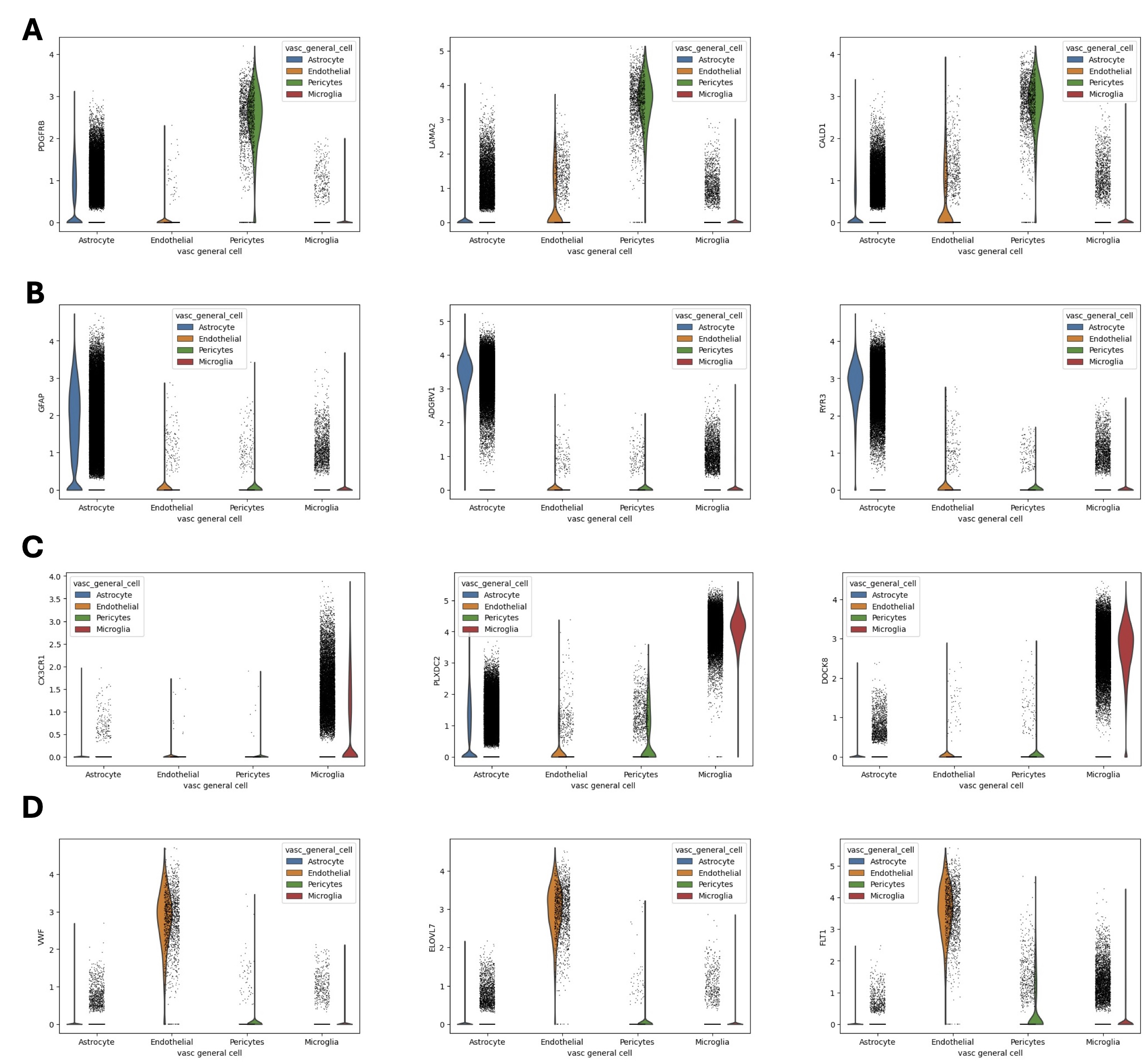

### Supplemental Figure 2

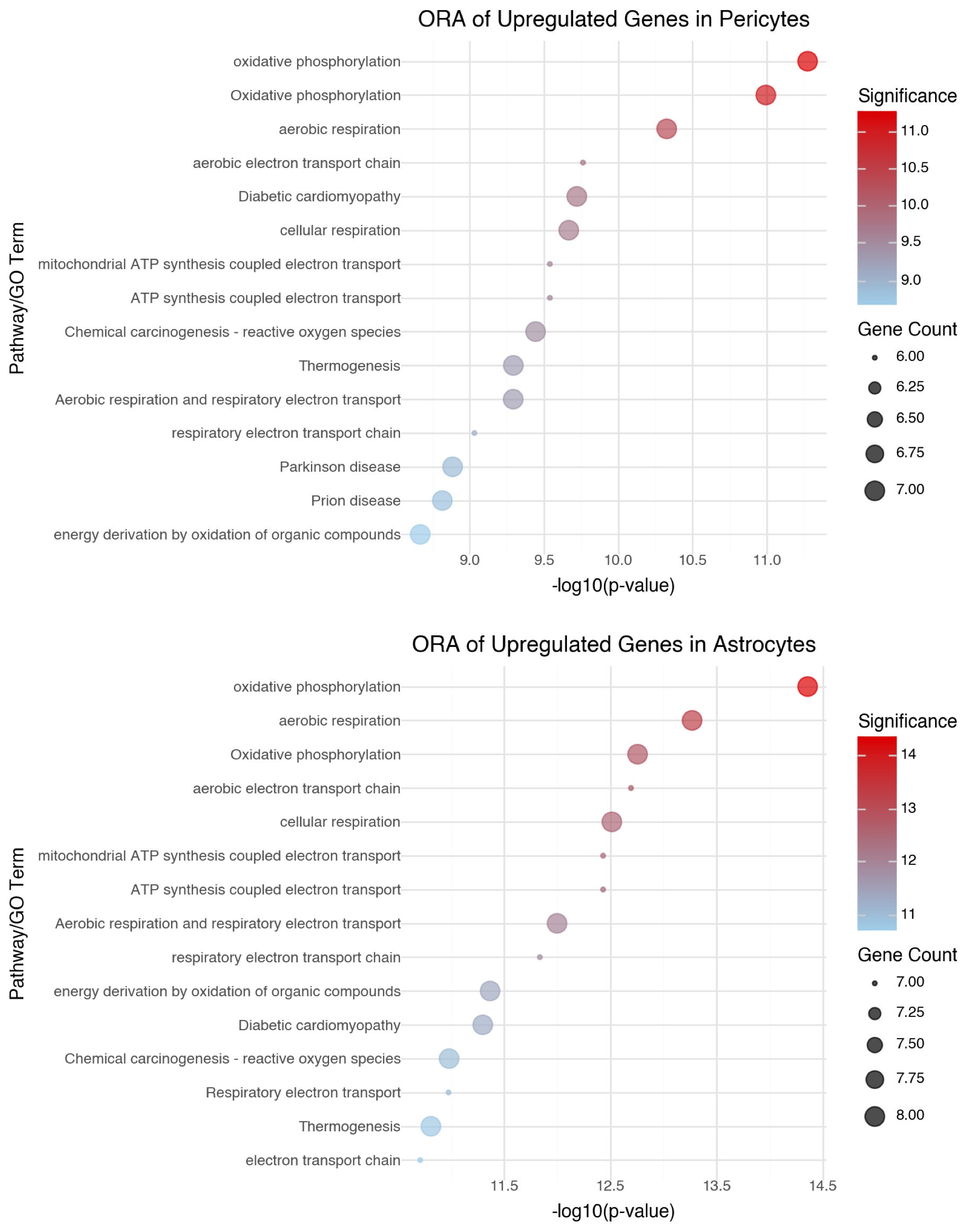

### Supplemental Figure 3

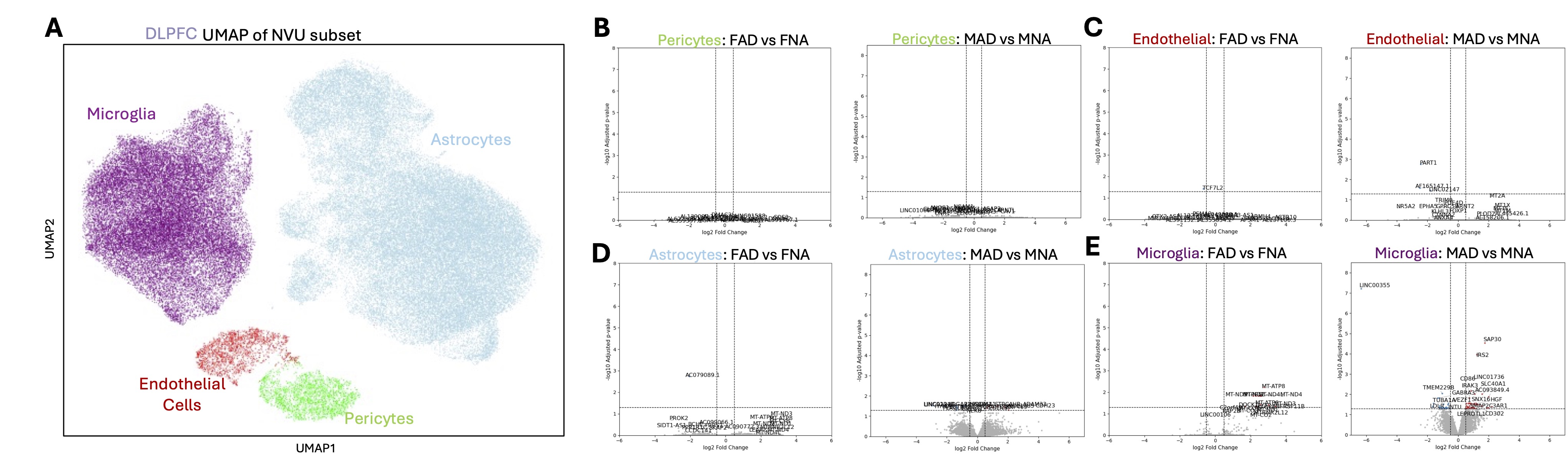

### Supplemental Figure 4

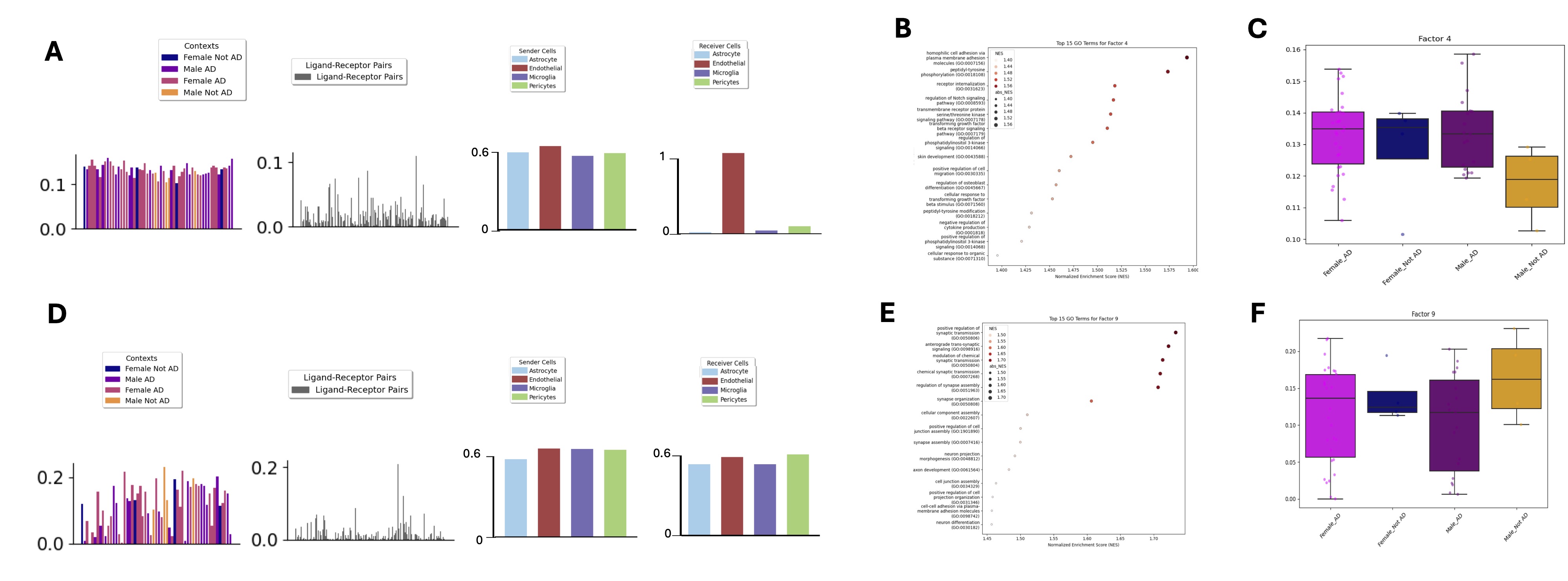
